# Roles of developmentally regulated KIF2A alternative isoforms in cortical neuron migration and differentiation

**DOI:** 10.1101/2020.05.07.081216

**Authors:** Cansu Akkaya, Dila Atak, Altug Kamacioglu, Busra Aytul Akarlar, Gokhan Guner, Efil Bayam, Ali Cihan Taskin, Nurhan Ozlu, Gulayse Ince-Dunn

## Abstract

KIF2A is a microtubule-depolymerizing kinesin motor protein with essential roles in neural progenitor division and axonal pruning during brain development. KIF2A is alternatively spliced in nervous tissue by specific RNA-binding proteins. However, how different KIF2A isoforms function during development of the cerebral cortex is not known. Here, we focus on three *Kif2a* isoforms expressed in mouse embryonic and postnatal cerebral cortex. We show that KIF2A is essential for dendritic pruning of primary cortical neurons in mice and that the functions of all three isoforms are sufficient for this process. Interestingly, only two of the isoforms can sustain radial migration of cortical neurons while a third isoform, lacking a key stretch of twenty amino acids, is ineffective. By proximity-labeling-based interactome mapping for individual KIF2A isoforms, we provide novel insight into how isoform specific interactions can confer changes to KIF2A protein function. Our interactome mapping identifies previously known KIF2A interactors, proteins localized to the mitotic spindle poles, and unexpectedly, also translation factors, ribonucleoproteins and proteins that are targeted to the mitochondria and ER, suggesting a novel transport function for KIF2A.

## Introduction

Alternative splicing (AS) is one of the major molecular processes underlying increased transcriptome and proteome diversity and associated organismal complexity in metazoans ^1-4^. High-throughput sequencing efforts have revealed that approximately 95% of all genes in the human body are alternatively spliced, generating more than 100,000 different mRNA isoforms, and the vast majority of isoform functions are completely unknown. AS is especially prevalent in the nervous system ^5,6^ and its impairment is often associated with neurological disease ^7,8^. AS is dynamically regulated during brain development, where it modulates important processes including, neural progenitor cell-to-neuron transition, migration, axon guidance, and synaptogenesis ^2,9,10^. Despite important advances in our understanding of how AS is regulated in a tissue-specific manner to allow transcriptome and proteome diversification, little is known about how individual alternative isoforms function to regulate nervous system development. In particular, only a small fraction of neurodevelopmental studies have investigated functions of distinct isoforms, and only the most abundant isoform is typically studied. Here, we have focused on the separable functions of three different isoforms of the Kinesin motor protein 2A (KIF2A) within the context of mouse cerebral cortex development.

KIF2A, a MT (microtubule)-depolymerizing kinesin, functions in mitotic spindle assembly ^11^ and primary cilium disassembly in mitotic cells ^12,13^ and in axonal pruning in differentiating neurons ^14,15^. Recent studies have suggested that KIF2A alternative splicing is regulated by neuron-specific RNA-binding proteins ^16^ and its isoform abundance dynamically shifts during the neural progenitor-to-neuron transition during development of the cerebral cortex ^3^. However, how the individual KIF2A isoforms function during cortical development has not been previously studied. We show that KIF2A is essential for proper dendritic pruning of cortical neurons and that all three KIF2A isoforms are interchangeable during this process. Moreover, we demonstrate that only two of the three KIF2A isoforms can functionally support radial neuronal migration in mice. We also reveal isoform-specific interacting protein partners of KIF2A by a proximity-ligation-based proteomic study carried out in a neuronal cell line. In addition to a number of MT-binding proteins, our interactome screen also identifies mitochondrial proteins and nuclear RNA-binding proteins as either common or isoform-specific interactors, pointing to a potential role for KIF2A in mitochondrial assembly and RNA localization. Our results highlight the importance of investigating isoform specific functions of gene products regulated by AS.

## Results

### *Kif2a* alternative isoform expression is developmentally regulated in mouse cerebral cortex

A previous study aimed at identifying the regulated targets of nElavl RNABPs (neuronal Elav-like RNA binding proteins) in the developing mouse cortex had identified *Kif2a* pre-mRNA as a top target ^16^. In mice the *Kif2a* gene consists of 20 coding exons, and nElavl proteins promote the inclusion of exon 18, which encodes for 37 amino acids rich in serine residues. While cloning these two major isoforms from the postnatal mouse cerebral cortex, we also identified a third isoform that includes exon 18, along with a shorter (by 20 amino acids) exon 5 generated as a result of alternative 5’ splice site selection (Fig.1A). We numbered these isoforms as *Kif2a.1* (exon 18+ and long exon 5), *Kif2a.2* (exon 18- and long exon 5) and *Kif2a.3* (exon 18+ and short exon 5) (Fig. 1A).

**Figure 1.**
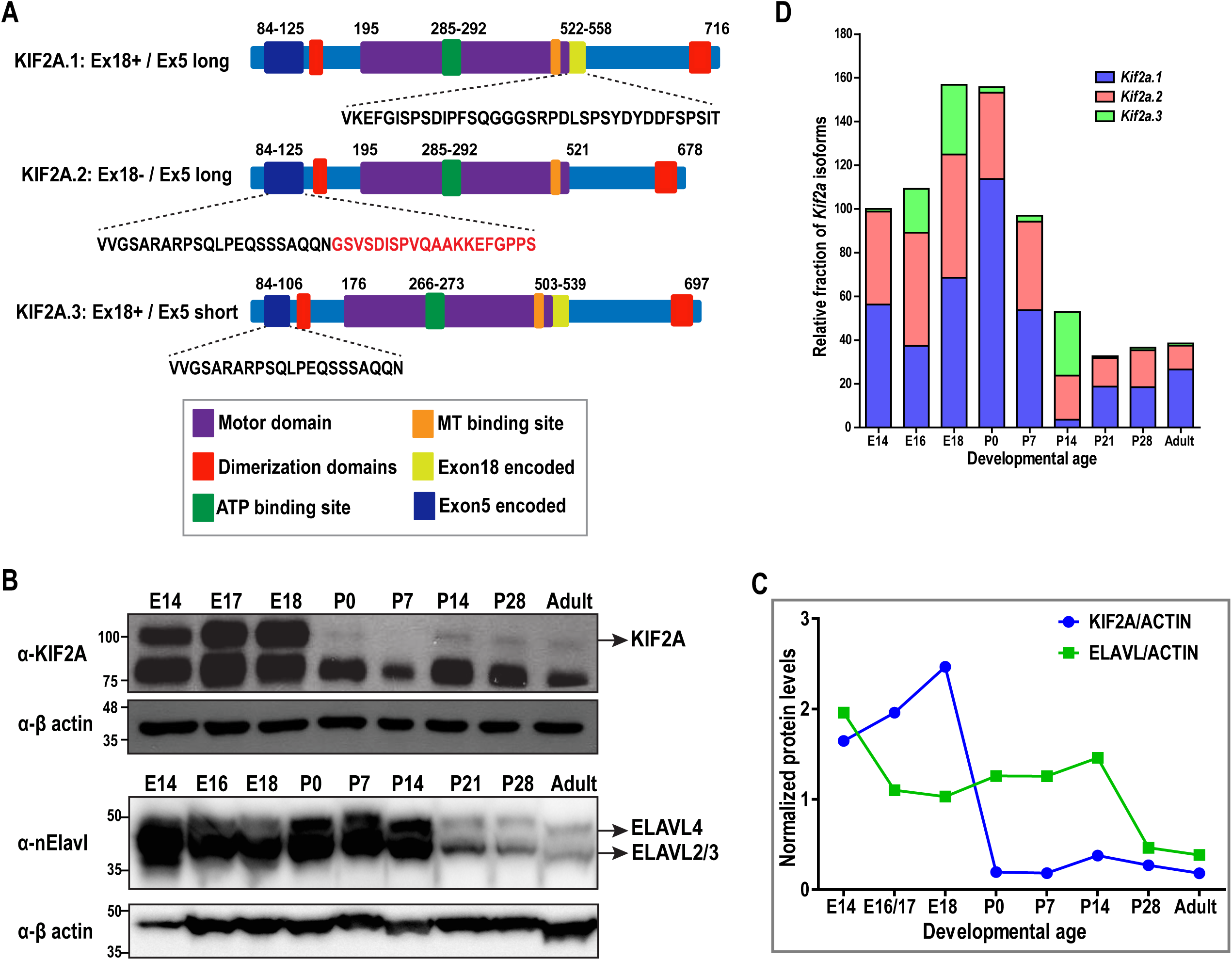
KIF2A expression is developmentally controlled in mouse cortex. **A**. Schematic of three different KIF2A isoforms shown as KIF2A.1, KIF2A.2 and KIF2A.3. Amino acid sequences of Exon 18, Exon 5-long and Exon 5-short are indicated and the amino acid sequence absent in Exon 5 of KIF2A.3 is highlighted in red. **B**. Total KIF2A (top) and nElavl (bottom) protein levels across development in mouse cerebral cortex. **C**. Quantification of immunoblot represented in (B). KIF2A and nElavl protein levels are normalized to B-ACTIN. **D**. The relative fraction of *Kif2a* isoform expression (normalized to first *Gapdh* and the levels measured in E14 tissue) quantified by qRT-PCR analysis. Data represents 3 biological x 2 technical replicates.

First, we tested the developmental expression patterns of these three *Kif2a* isoforms at the mRNA level, along with overall protein abundance of KIF2A and nElavl RNABPs. Since an antibody specific to individual isoforms of KIF2A proteins does not exist, and generating isoform specific antibodies is exceedingly challenging, we were able to measure only total KIF2A protein levels. We found the expression dynamics of nElavl and KIF2A protein abundance to correlate and generally exhibit elevated expression levels in embryonic cortex and to be downregulated postnatally (Fig.1B and C), suggesting that nElavl RNABP might regulate *Kif2a* mRNA stability in addition to its AS. Next we quantified the relative mRNA abundance of the three *Kif2a* isoforms in the developing cerebral cortex by quantitative RT-PCR. Our results demonstrated that *Kif2a.1* and *Kif2a.2* were the predominant isoforms, manifesting higher embryonic expression levels when compared to postnatal and adult stages. *Kif2a.3* was generally the minor isoform, with expression peaking during late embryogenesis and once again during the second postnatal week (Fig.1D). Taken together, our results indicate an overall trend in which expression of KIF2A isoforms is high during embryonic stages of the developing mouse cerebral cortex, a time period when neural progenitor proliferation and active neuronal migration ensues.

### KIF2A isoforms are interchangeable in their roles for dendritic arborization of cortical neurons

To further understand whether the three KIF2A isoforms may have divergent functions we asked whether AS had an effect on their subcellular localization. For this purpose, we imaged all three KIF2A isoforms in Neuro2A cells and primary cortical neurons after expressing KIF2A-EGFP fusion proteins of individual isoforms. Our images revealed that all three KIF2A isoforms are localized predominantly to the cytoplasm and nucleus in both cell types (Fig.2A and Suppl. Fig.1). Furthermore, we detected axonal and dendritic localization in cortical neurons for all isoforms (Fig.2A). Hence, our results suggest that alternative splicing of KIF2A does not influence its subcellular localization in cultured neurons.

**Figure 2.**
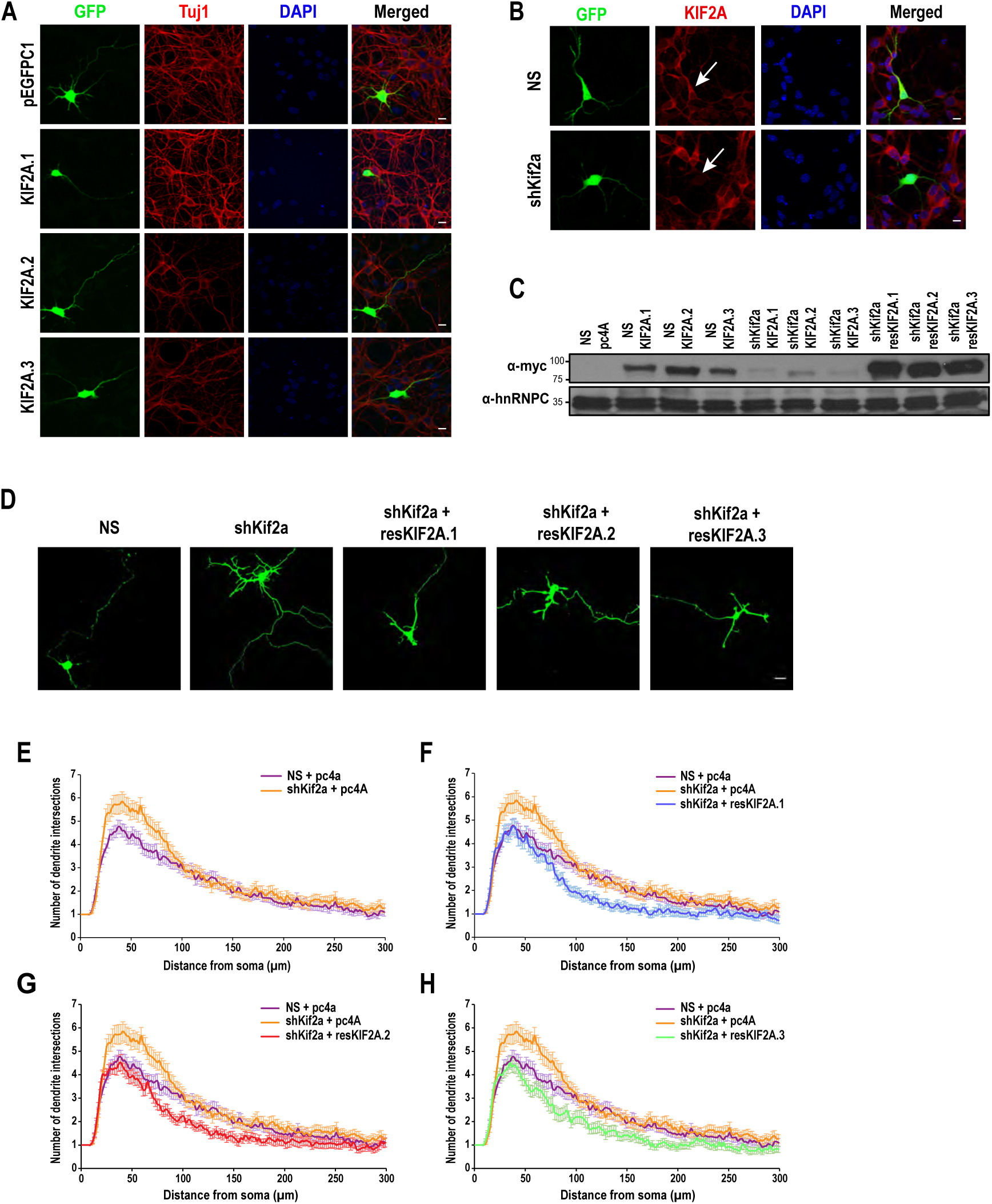
KIF2A isoforms function in dendritic pruning. **A**. Primary cortical neurons from embryonic day 18.5 (E18.5) embryos transfected with plasmids expressing pEGFPC1-KIF2A (KIF2A.1, KIF2A.2 and KIF2A.3 separately) at 2 day *in vitro* (DIV), fixed and immunostained with anti-GFP at 4 DIV. Microtubules were visualized with anti-Tuj1 antibody (red). **B**. E14.5 primary cortical neurons were transfected with non-silencing shRNA (NS) or shRNA against KIF2A (shKif2a) at 2 DIV, fixed at 4 DIV and immunostained against GFP (green) and KIF2A (red) proteins. Transfected cells were identified based on their co-expression of GFP from NS (non-silencing) or shRNA expressing plasmid. White arrows indicate transfected neurons. **C**. Immunoblotting against myc tagged *Kif2a* isoforms, whose expression is suppressed by co-expression of *shKif2a*. The *resKif2a* cDNAs are resistant to silencing by *shKif2a*. hnRNPC is used as a loading control. **D**. Representative images of primary cortical neurons from E14.5 embryos co-transfected with NS or *shKif2a* along with either pcDNA4A backbone or resKIF2A at 2 DIV, fixed at 4 DIV and imaged with confocal microscopy. **E, F, G and H**. Dendrite arborization was quantified by Sholl analysis ^19,65^. ((E), NS-shKif2a; (F), NS-shKif2a-resKIF2A.1; (G), NS-shKif2a-resKIF2A.2; and (H), NS-shKif2a-resKIF2A.3). n=60 for each condition derived from two separate neuronal cultures. Bars represent S.E.M. Unpaired two-tailed *t* test, p value: * p<0.05, **p<0.005. Scale bar in A, B and D, 10 µm.

Previous studies had demonstrated a role for KIF2A in pruning of axon collaterals of cortical neurons and sensory neurons in mice ^15,17^. Since regulation of MT dynamics is also critical for dendritic arbor growth and pruning ^18^, we next evaluated the role of KIF2A isoforms in dendrite development. Toward this aim, we knocked down *Kif2a* expression in primary cortical neurons by expressing a shRNA targeting a common sequence in all three isoforms (Fig.2B). We co-expressed soluble EGFP for imaging neuronal processes and measured dendritic arborization by Sholl analysis, which is a quantitative measure of dendritic length and branching (Fig.2D) ^19^. Our results revealed a significant increase in the number of primary dendrites, suggesting diminished capacity for MT depolymerization and dendritic pruning in the absence of KIF2A (Fig.2E). In order to test the ability of different KIF2A isoforms to carry out its role in dendritic pruning, we knocked down all *Kif2a* expression, and then returned one isoform at a time by re-introducing shRNA-resistant forms of individual isoforms (*resKif2a.1, resKif2a.2* and *resKif2a.3*) (Fig.2C and D). As above, we quantified dendritic development by Sholl analysis. Our analysis revealed that all three isoforms of KIF2A were able to suppress the growth of superfluous dendrite branches and could reverse the KIF2A loss-of-function phenotype equally well (Fig.2F, G and H). Taken together, we conclude that KIF2A is required for normal dendrite development in cortical neurons and all three KIF2A isoforms may carry out this function.

### Specific KIF2A isoforms appear to play a role in radial migration of cortical neurons

Previous studies have demonstrated that KIF2A is required for cortical neuronal migration. Mutations in *Kif2a* affect neuronal positioning in the cerebral cortex and cause cortical malformations in humans ^20-22^. Consequently, we investigated whether KIF2A’s role in cortical neuronal migration is isoform-dependent. During development, the neural progenitor cells divide in the ventricular zone (VZ) of the embryonic telencephalon. After exiting the cell cycle, the early neurons radially migrate through the subventricular and intermediate zones (SVZ-IZ) until they reach the most superficial layer of the developing cortical plate (CP) ^23^. In order to assess the role of different KIF2A isoforms in this process, we knocked down expression of all *Kif2a* isoforms, then re-introduced one isoform at a time by expressing shRNA-resistant *Kif2a* cDNAs in ventricular neural progenitors of embryonic day 14.5 (E14.5) cerebral cortex by *in utero* electroporation. We also electroporated a plasmid expressing the tdTomato marker and quantified the extent of neuronal migration by quantifying the number of tdTomato-positive neurons as a function of distance away from the ventricular zone (Fig.3A and B). As expected, approximately 45% of tdTomato-positive neurons had reached the CP, 50% were migrating along the SVZ-IZ region and 5% were localized to the VZ in control samples at three days post-electroporation (Fig.3A). Consistent with previous observations ^20,21^, there was a dramatic reduction in the relative fraction of tdTomato-positive neurons localized to the CP in *Kif2a* knockdown samples, and deficient migration caused most of the neurons to accumulate in the SVZ-IZ (Fig.3A). In cortices where *Kif2a* expression was rescued with either *resKIF2A.1* or *resKIF2A.2*, neurons could exit the SVZ-IZ and reach the CP, at levels comparable to control samples (Fig.3C, D, F and G). In contrast, *resKIF2A.3*-expressing neurons exhibited a severe migration defect, and the number of neurons that could migrate to the cortical plate was significantly reduced (Fig.3E and H). It is noteworthy to mention that KIF2A.3 differs from the other two isoforms by 20 amino acids encoded by exon5, which corresponds to a disordered protein region predicted by the IUPred2A prediction tool ^24^ (Suppl. Fig.2). Interestingly, tissue specific splicing of disordered protein regions is increasingly acknowledged as a means to increase interaction networks across tissue ^25^. Taken together, our results suggest that KIF2A.1 and KIF2A.2 isoforms, but not the KIF2A.3 isoform, can support neuronal migration in the developing cortex.

**Figure 3.**
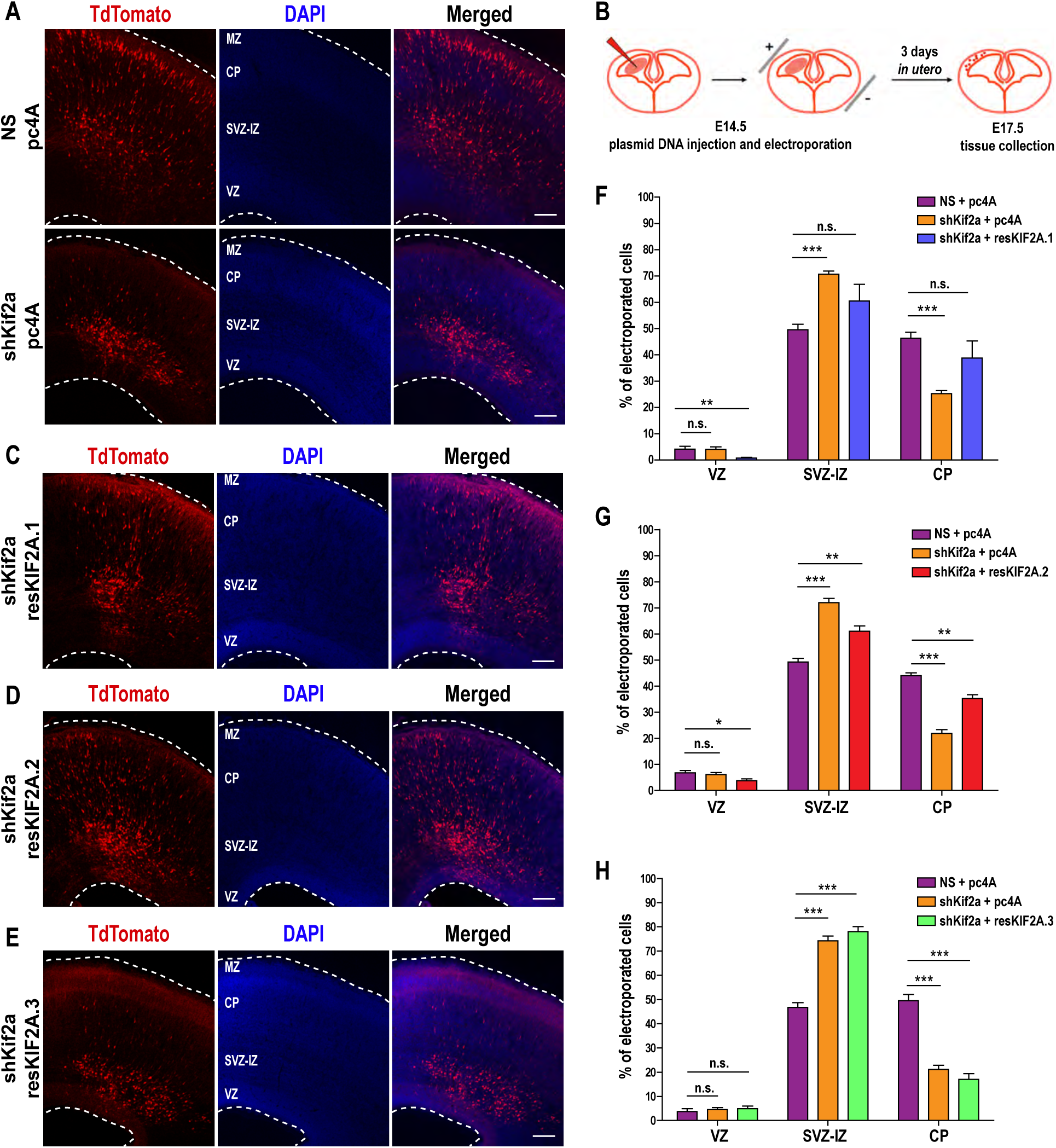
Role of different KIF2A isoforms in radial migration. **A**. DAPI staining of E17.5 coronal sections *in utero* electroporated with either mCherry-tagged NS or shKif2a along with pcDNA4A (pc4A) and tdTomato at E14.5. **B**. Relevant plasmid DNA expressing *shKif2a, resKif2a* and *tdTomato* were injected into the ventricle of E14.5 mice. Following a brief pulse of electroporation embryos were replaced back into the uterus and allowed to develop for 3 days *in utero*. At E17.5 the embryonic cortices were collected for migration analysis. **C, D and E**. Embryos electroporated *in utero* with mCherry-tagged *shKif2a* along with one of the *resKif2a* (resKIF2A.1 (C), resKIF2A.2 (D) or resKIF2A.3 (E)) and tdTomato at E14.5. Boundaries of cortical sections were indicated as white dashed lines. MZ, marginal zone; CP, cortical plate; SVZ-IZ, subventricular zone-intermediate zone; VZ, ventricular zone. **F, G and H**. Quantification of *in utero* electroporation experiment of resKIF2A.1 (F), resKIF2A.2 (G) and resKIF2A.3 (H). Cortical sections were divided into zones. Percentages of tdTomato-positive neurons counted in individual zones are plotted. (For KIF2A.1 a total of n=4017 for NS, n=4572 for shKIF2A, n=2696 for resKIF2A.1 from 3 biological replicates; for KIF2A.2 a total of n=2477 for NS, n=3160 for shKIF2A, n=3558 for resKIF2A.2 from 2 biological replicates; for KIF2A.3 a total of n=3189 for NS, n=4456 for shKIF2A, n=3549 for resKIF2A.3 from 3 biological replicates). Data is represented as bar graphs. Lines represent S.E.M. Unpaired two-tailed *t* test, p value: *p<0.05, **p<0.005, ***p<0.0001. Scale bar in A, C, D and E, 100 µm.

### Proximity-labeling proteomics analysis of KIF2A isoforms reveal novel isoform-specific interactors

Experimental analysis and computational modeling have suggested that an important function of alternative splicing is to increase phenotypic complexity by expanding the interactome ^25-27^. Therefore, we next asked whether different KIF2A isoforms were indeed characterized by different protein interaction networks. Toward this goal, we performed a proximity-labeling method which allows the identification of interactors of a protein of interest by its expression as a promiscuous biotin-ligase (the BirA* gene) fusion in the cell type of choice, followed by streptavidin based affinity purification and mass spectrometry analysis (BioID) ^28^. In order to harvest sufficient total biotinylated protein, we decided to utilize the easily transfectable Neuro2A neuronal cell line for our BioID experiments. Initially, we confirmed that our KIF2A-BirA* fusion proteins were functional. For this, we transiently transfected KIF2A-BirA* fusions for each of the three isoforms, supplemented the cells with 0.05mM biotin, and confirmed that indeed biotinylated proteins co-localized with expressed KIF2A-BirA* fusion proteins (Fig.4A). Next, we verified that biotinylation was dependent upon the external supply of biotin, and that the background signal in the absence of external biotin following streptavidin pulldown was very low (Fig.4B and C).

**Figure 4.**
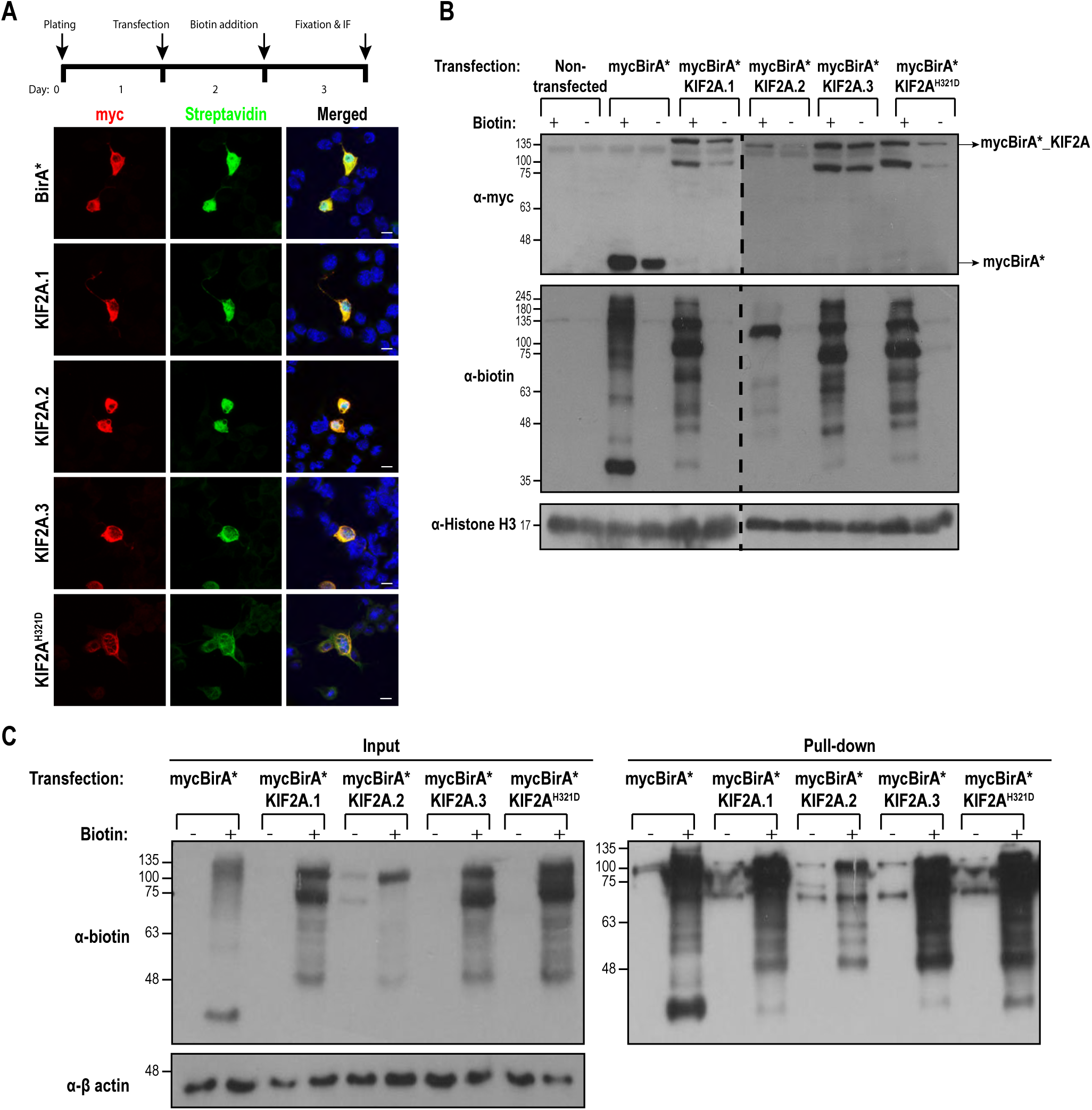
Proximity biotin ligation assay of KIF2A isoforms in Neuro2A cells. **A**. Immunofluorescence staining of myc (red), Streptavidin (green) and DNA (blue) in Neuro2A cells transfected with myc*BirA\**, myc*BirA*-Kif2a.1, mycBirA*-Kif2a.2, mycBirA*-Kif2a.3* or *mycBirA*-Kif2a*^*H321D*^ separately. Transfected cells were identified based on myc immunostaining. Scale bar, 10 µm. **B**. Top panel: Immunoblotting with anti-myc showing expression of mycBirA*-KIF2A in Neuro2A cells. Middle panel: Immunoblotting with anti-biotin confirming biotinylation. Bottom panel: Anti-Histone H3 as loading control. **C**. Pull-down with streptavidin beads followed by anti-biotin immunoblotting of Neuro2A cell lysates. Biotin supplementation to cell media is indicated.

Having validated the BioID approach, we scaled up our protocol and deployed tandem mass spectrometry for analysis of purified proteins after streptavidin pulldown. We analyzed three biological replicates for each isoform, and based on PSM (peptide spectra match) values, we filtered out proteins identified in only one of the replicates, those with FDR<0.05, and those with fold-change<1.5. Non-transfected Neuro2A cells were used as control for background subtraction. Following these cut-off criteria, we identified a total of 63 potential interactors for KIF2A.1, 37 for KIF2A.2, and 40 for KIF2A.3 (Fig.5A, G and Suppl.Table.1). As expected, KIF2A itself was among the highest ranking hits for each of the three isoforms. We identified proteins that were common among all three KIF2A isoforms, shared in all possible pairwise combinations, as well as those that were unique for KIF2A.1, KIF2A.2 or KIF2A.3 (Fig.5A and G; Suppl.Table.1). Next we carried out gene ontology and interactome analysis using the String Database ^29-32^ separately with interactors of each of the KIF2A isoforms (Fig.5D, E and F). As expected, MT-associated proteins which function at the spindle poles during mitotic division were identified among the interactomes of all three isoforms (eg. CKAP5, NUMA1, CEP170 and MAP4). Our analysis revealed the highest number of interactors for KIF2A.1. Intriguingly, these were enriched for mitochondrial proteins with functions in mitochondrial gene expression (eg. TIMM44, PNPT1, LARS2, RARS2 and TSFM) and metabolism (eg. GLS, ALDH2, OAT, SUCLG2 and SUCLA2), nuclear RNA regulation (eg. SRSF3, SRSF4, SRRM2 and HNRNPA1) (Fig.5D and G). KIF2A.2 displayed the least number of interactors, many of which were shared with KIF2A.1, with similar functions in mitochondrial metabolism and nuclear RNA transport (Fig.5E and G). KIF2A.3 displayed the highest relative fraction of unique interactors. Its interactome was enriched in functions relating to nuclear RNA transport and regulation. Unlike the other two isoforms, KIF2A.3 was largely devoid in genes functioning in the mitochondria (Fig.5F and G). Intriguingly, all KIF2A isoforms also appeared to interact with different eukaryotic translation factors (EIF5, EIF4G2, EEF1B2 and EIF3E). In sum, our data identifies common and unique interactors of the three KIF2A isoforms.

**Figure 5.**
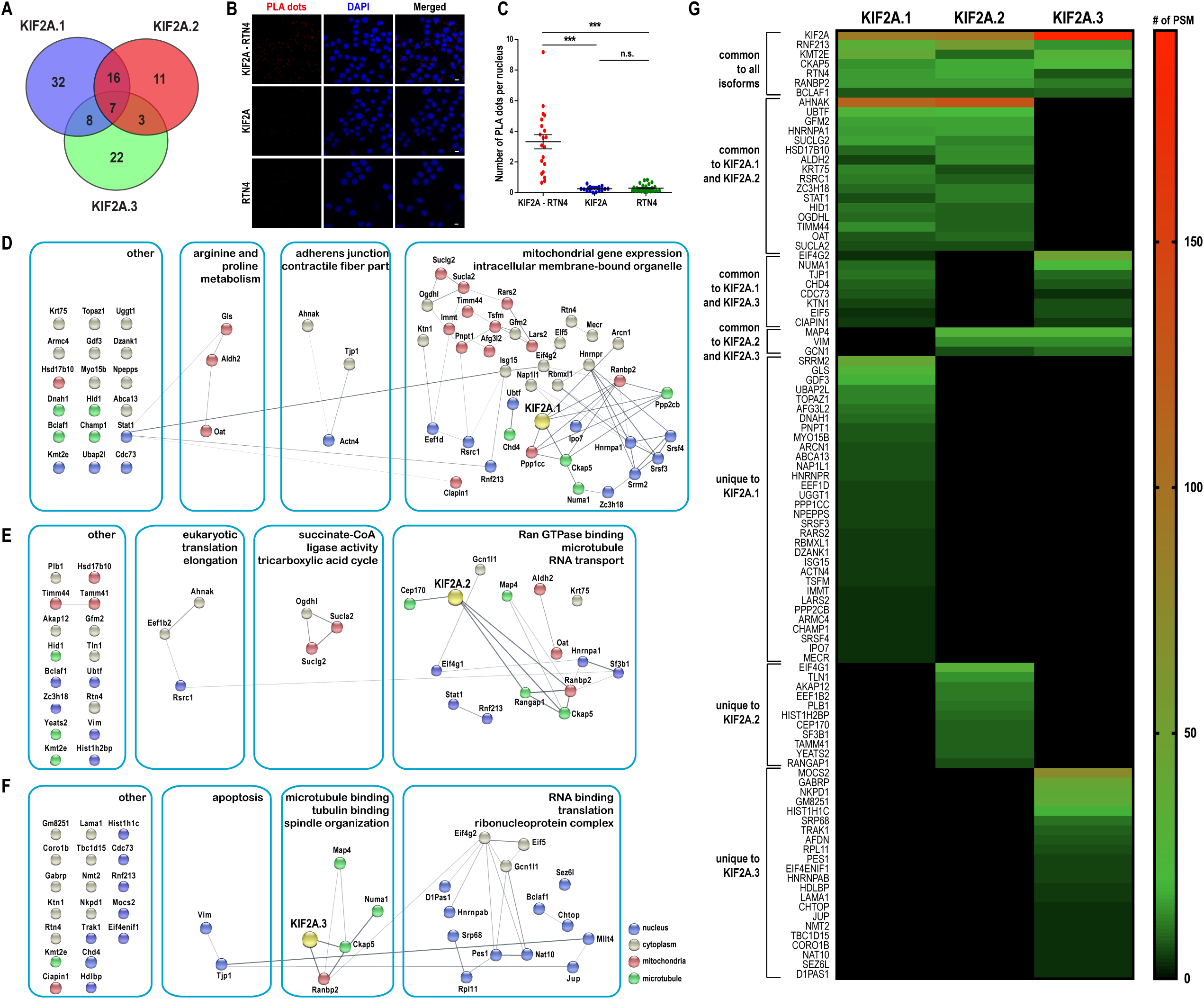
Protein interactome of KIF2A isoforms is revealed by a proximity ligation assay. **A**. *Venn* diagram represents the number of common and specific proximal protein partners of KIF2A isoforms. n=3 biological x 2 technical replicates. **B**. KIF2A interacts with RTN4 in a spatiotemporal resolution. Immunofluorescence staining of PLA dots (red), DNA (blue) in Neuro2A cells. Scale bar, 10 µm. **C**. Spatial analysis of KIF2A-RTN4 interaction by PLA assay. 20 image frames (n=25 frames for RTN4) chosen randomly. 3.2% paraformaldehyde (PFA) fixed-Neuro2A cells were incubated KIF2A-RTN4 antibody pairs (n=1052) for interaction analysis. Fixed cells incubated with only KIF2A (n=684) or only RTN4 (n=373) were used as control. Lines indicate the mean number of PLA dots per nucleus. Bars represent S.E.M. p value: ***p<0.0001 by one-way ANOVA with Bonferroni post hoc test. **D, E and F**. Interaction networks of significantly enriched proteins for KIF2A.1 (D), KIF2A.2 (E) and KIF2A.3 (F) are constructed by STRING database ^32^. Highly interconnected subnetworks are highlighted with blue rectangles and enriched GO and KEGG terms of each network are reported ^29,31^. The ones with more than one annotated location are only illustrated with one compartment. Isoforms are indicated as yellow circles. Proteins are colored according to their localization in the cell; gray, cytoplasm; blue, nucleus; green, microtubule; and, red, mitochondria. Gray lines represent the known interactions from STRING database and thickness indicates an increase in interaction confidence. **G**. Heat map generated through PSM (peptide spectra match) values of proteins that were identified with BioID method for three KIF2A isoforms.

Next we decided to focus upon one of the interacting polypeptides. Specifically, we decided to investigate interactions between endogenous KIF2A and RTN4 (Reticulon-4 or also known as NOGO-A), which we identified in all three KIF2A isoform samples. RTN4 localizes to the ER membrane and to the plasma membrane. In the ER it has been show to regulate vesicular transport and structural stabilization of the ER network ^33^. In the nervous system RTN4 acts as is an inhibitor of neurite outgrowth; it inhibits migration and sprouting of central nervous system (CNS) endothelial (tip) cells and axons by providing inhibitory signals ^34-36^. Our co-immunofluorescence staining with KIF2A and RTN4 antibodies displayed a partial co-localization (Suppl.Fig.3). Next, we confirmed the interaction between KIF2A and RTN4 by proximity ligation assay (PLA), which is a sensitive and specific method for detecting protein interactions with spatial resolution ^37^. Our PLA results revealed a significant interaction between KIF2A and RTN4, compared to negative controls where only anti-KIF2A or anti-RTN4 primary antibodies were used for the PLA protocol (Fig.5B and C).

### KIF2A disease mutation causes rewiring of its protein interactome

Having established a system to uncover the protein interaction network of KIF2A, we next decided to turn our attention to KIF2A (KIF2A^S317N^ and KIF2A^H321D^) mutations that cause developmental cortical malformations in humans ^20,22^. Initially, we generated identical disease-associated point mutations at conserved residues in the backbone of mouse *Kif2a.1* cDNA. To evaluate how these two KIF2A.1 mutants localize subcellularly in neuronal cells, we expressed EGFP-KIF2A^S317N^ and EGFP-KIF2A^H321D^ in Neuro2A cells and mouse primary cortical neurons. Consistent with previous observations, both KIF2A^S317N^ and KIF2A^H321D^ mutants were sequestered along polymerized MTs and did not display the typical cytoplasmic and nuclear distribution pattern we observed for wild-type KIF2A.1 (Fig.6B and Suppl.Fig.4A).

**Figure 6.**
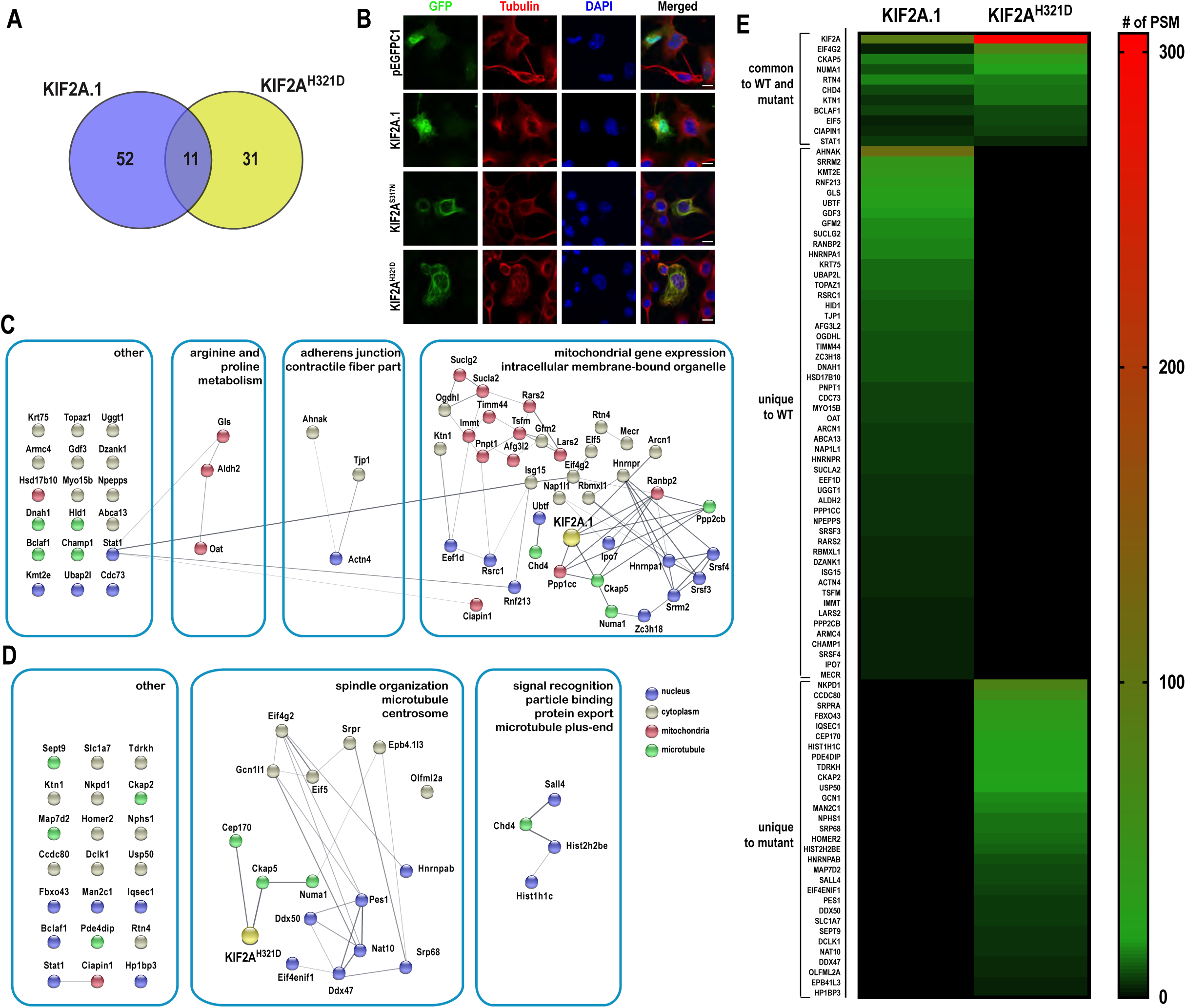
Protein interactome of KIF2A^H321D^ is revealed by a proximity ligation assay. **A**. *Venn* diagram representing the number of common and specific proximal protein partners of KIF2A.1 and KIF2A^H321D^. n=3 biological x 2 technical replicates. **B**. Both KIF2A^S317N^ and KIF2A^H321D^ are co-localized predominantly to microtubules in Neuro2A cells. Neuro2A cells were transfected with plasmids expressing pEGFPC1-tagged KIF2A.1, KIF2A^S317N^ or KIF2A^H321D^ and immunostained with anti-GFP (green) or anti-α-Tubulin (red) antibodies. Scale bar 10 µm **C and D**. STRING interaction ^32^ network of candidate proteins identified for KIF2A.1 (C) and KIF2A^H321D^ (D) ^29,31^. Gray Lines indicate known interactions and thickness represent confidence of interaction. **E**. Heat map created based on the PSM values of proteins that are identified with BioID of KIF2A.1 and KIF2A^H321D^.

Next, we analyzed results from our previous BioID proximity labeling experiments in which KIF2A^H321D^ (along with the three isoforms) was used as a bait. We compared proteins identified in KIF2A^H321D^ and KIF2A.1 samples as the mutation was created within this isoform. Our comparison revealed widespread rewiring of the interactome upon introduction of the H321D mutation. We identified 52 proteins that were unique to KIF2A.1, 31 unique to KIF2A^H321D^ and 11 that were common to both (Fig.4A-C and 6A and E; Suppl.Table.1). Next, we generated a gene ontology and interactome map for KIF2A^H321D^ and compared it to that of KIF2A.1, using the String database ^29-32^. The relative fraction of cytoplasmic, nuclear and MT-associated genes was preserved in KIF2A^H321D^ however, the identities were different (Fig.6C, D and E). The most striking observation was the depletion of mitochondrial protein from the interactome of KIF2A^H321D^ in comparison to KIF2A.1 (16 mitochondrial proteins out of a total of 63 proteins for KIF2A.1 and 1 out of 42 for KIF2A^H321D^).

Finally, we asked whether these dominant mutations affected dendrite development in cortical neurons. For this purpose, we co-transfected E14.5 mouse primary cortical neurons with pEGFPC1 along with either one of the *Kif2a* mutants (*Kif2a*^*S317N*^ or *Kif2a*^*H321D*^) and carried out Sholl analysis. We observed dendritic arbor development comparable between cultures transfected with control cultures and cultures expressing mutant KIF2A proteins (Suppl.Fig.4B and C). Unexpectedly, our results suggested that disease-causing dominant mutations in *Kif2a* may not affect dendrite development in primary cortical neurons.

## Discussion

*Kif2a* is an alternatively spliced gene, and isoform abundance is developmentally regulated ^3,16^. Here, we reveal that all three KIF2A isoforms can rescue a dendritic pruning defect caused by knockdown of all KIF2A expression in cortical neurons. However, only KIF2A.1 and KIF2A.2, but not KIF2A.3, are able to support migration of neurons in the developing cortex (Fig.7). KIF2A.3 differs from the other two isoforms by a 20 amino acid-long serine-rich disordered protein region located immediately N-terminal to its dimerization domain. Currently, we do not know how this short stretch of amino acids affects KIF2A function. However our results from our isoform-specific proximity-labeling screen demonstrated that the protein interactome of KIF2A.3 is different from the other two isoforms.

**Figure 7.**
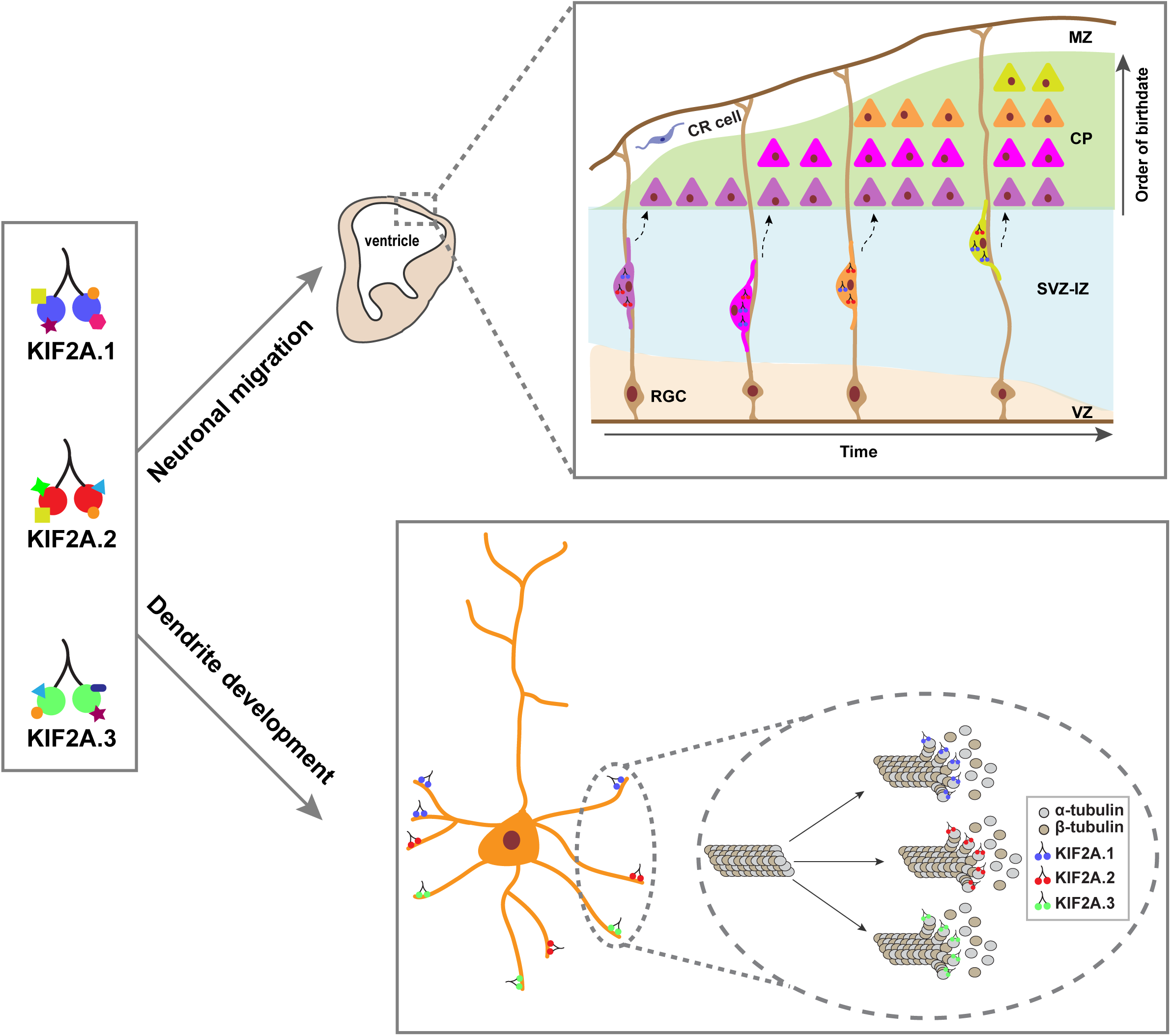
Model of KIF2A isoforms in cortical dendritic pruning and neuronal migration. Three alternatively spliced KIF2A isoforms (KIF2A.1 in blue, KIF2A.2 in red and KIF2A.3 in green) have unique and common interactors. Upper panel shows how neurons migrate radially during cortical development in an “inside-out” manner and that KIF2A’s role in cortical neuronal migration is isoform dependent. Only KIF2A.1 and KIF2A.2 but not KIF2A.3 can support the radial neuronal migration in developing cortex. Lower panel summarizes role of KIF2A isoforms in dendrite development of primary cortical neurons in mice. KIF2A is essential for dendritic pruning and all three KIF2A isoforms can fulfill this function by depolymerizing microtubules. RGC, radial glial cell; CR, Cajal-Retzius; MZ, marginal zone; CP, cortical plate; SVZ-IZ, subventricular zone-intermediate zone; VZ, ventricular zone.

Our BioID assay carried out in a neuronal cell line identifies a number of centrosomal proteins with roles in bipolar spindle assembly, of which NUMA and CEP170 are known KIF2A interactors ^13,38,39^. In mitotic cells KIF2A is essential for bipolar spindle assembly and at the poles it depolymerizes minus end MTs to contribute to poleward chromosome movement ^11,40,41^. Here, we provide additional candidate KIF2A interacting proteins such as KMT2E, CKAP5, RANBP2 and CHD4, which also localize to bipolar spindles. Further investigations on how KIF2A and these candidate proteins functionally interact is likely to provide new mechanistic understanding of bipolar spindle assembly and chromosome segregation.

Previous findings had demonstrated an essential role for KIF2A in axonal pruning ^15,17^, neuronal migration ^17,42^ and in primary cilium disassembly during mitotic division of neural progenitors ^12,13,20,21^. Since Neuro2A cells are actively dividing and do not possess primary cilia, our BioID assay did not identify primary cilium specific proteins. However, several of the candidate KIF2A interactors are likely to have important functions during development of the cerebral cortex. For example, we have verified interactions between the neurite outgrowth inhibitor RTN4/NOGO-A and endogenous KIF2A in Neuro2A cells. Interestingly, RTN4/NOGO-A acts a negative regulator of radial neuron migration and cortical lamination is defective in *Rnt4* knockout mice ^43^. If and how KIF2A-RTN4 interactions affect radial neuron migration is currently a wide-open question. Among other potential interactors is CEP170, a KIF2A.2 specific interactor, which regulates primary cilia disassembly and neural progenitor division in the ventricular zone ^13^. CHD4 is a candidate interactor of KIF2A.1 and KIF2A.3 isoforms, and a chromatin remodeling histone deacetylase, promoting the proliferation of neural progenitors at the cortical ventricular zone during development ^44^. Finally, mutations in KMT2E, a lysine methyl transferase localized to spindle poles and a candidate interactor of all KIF2A isoforms, result in a range of neurodevelopmental disorders in humans, often manifesting with microcephaly and epilepsy ^45^. Further characterization of these potential KIF2A interactors will reveal important details of how KIF2A controls early cortical development.

We also identify unexpected KIF2A interactors such as a number of mitochondrial- or ER-targeted proteins, RNA processing enzymes, and translation factors. We recognize that a subset of these hits may represent potential contaminants in our dataset. However, given that many of them display clear isoform-specificity we wish to speculate on the exciting implications of these findings. Several of these proteins have well-described roles in MT-based cargo trafficking. For example, KTN1 and TRAK1 are both kinesin adaptor proteins with essential roles in mitochondrial transport along MTs ^46-50^. Paradoxically, mitochondria and ER subcellular distribution is not affected in *Kif2a* knockout cells ^17^ raising the possibility of genetic redundancy among different kinesins during organelle trafficking.

Here we have demonstrated that the KIF2A.3 isoform cannot support neuronal migration. In addition, previous studies have shown that the KIF2A^H321D^ mutation is causal for migration defects in humans and mice ^20,22^. Interestingly, we have noticed that the interactomes of KIF2A.3 and KIF2A^H321D^ display a considerable number of common targets not identified in our KIF2A.1 and KIF2A.2 interactome datasets. Many of these hits are known or suggested components of ribonucleoprotein complexes (HNRNPAB, EIF4ENIF1, PES1 and NAT10) ^51-55^. Furthermore, KIF2A.3 and KIF2A^H321D^ datasets are largely devoid of much of the mitochondrial proteins identified in interactome datasets of KIF2A.1 and KIF2A.2. It is noteworthy to mention that several studies have demonstrated MT- and kinesin-dependent transport mechanisms for different kinds of RNA-protein complexes, including mRNAs that are targeted to the mitochondria ^56-59^. Whether KIF2A plays a direct role in MT-based RNA-protein complex transport, or might generally be in close proximity to a number of MT-associated-cargo due to its function in MT depolymerization, remains to be discovered. While further analysis is essential to resolve the biological relevance of our findings, our results showcase the utility of our approach to identify potential interactors of different KIF2A isoforms with relevance to pathogenesis of human cortical malformations.

## Material and Methods

### Cloning

Mouse *Kif2a.1, Kif2a.2* and *Kif2a.3* and *Kif2a*^*H321D*^ were subcloned from the pEGFPC1 vector into *Xho*I and *EcoR*I sites of the pcDNA3.1(-)-mycBirA backbone vector. (BirA*, N terminal; kindly provided by Elif N. Firat-Karalar, Koç University, Istanbul). The primer sequences were as follows: Kif2a_BirAN_F (aaactcgagatggtaacatctttaaatgaagat); Kif2a_BirAN_R (aaagaattcttagagggctggggcctcttggg). H321D and S317N point mutations were created by site-directed mutagenesis of mouse pEGFPC1-Kif2a.1. The primer sequences were as follows: Kif2a_c959G>A_F (gcagactggaaatgggaaaactcatactatgg); Kif2a_c961C>G_F (gcagactggaagtgggaaaactgatactatgg); Kif2a_sdm_R (ccataagcaaagcacgtagccatgccccttt). Non-silencing (NS) and shKif2a short hairpin RNAs (shRNA) were cloned into pSUPER-neo-EGFP (www.oligoengine.com) as described in the manual. For *in utero* electroporation experiments, the EGFP cassette was exchanged with an mCherry cDNA by subcloning into the *Age*I and *Not*I sites of pSUPER-neo. Three isoforms of shKif2a-resistant cDNA (resKIF2A.1, resKIF2A.2 and resKIF2A.3) were created by site-directed mutagenesis in pcDNA4A backbone vector by introducing three silent mutations (A1521T, C1524G, T1527C) within the shKif2a target sequence (ggaatggcatcctgtgaaa) ^60^. The primer sequences were as follows: NS_F(gatccccgcgcgatagcgctaataatttttcaagagaaaattattagcgctatcgcgcttttta); NS_R(agtctaaaaagcgcgatagcgctaataattttctcttgaaaaattattagcgctatcgcgcggg);shKif2a_F(gatccccg gaatggcatcctgtgaaattcaagagatttcacaggatgccattccttttta);shKif2a_R(agctaaaaaggaatggcatcctgtgaa atctcttgaatttcacaggatgccattccggg);resKIF2A_F(ccaggaatggcttcgtgcgaaaatactct);resKIF2A_R(a gagattgtggcaatcatgcaggtacgag). All constructs were confirmed by Sanger sequencing.

### Primary cortical cultures

Primary cortical cultures were prepared as previously described ^61^. Briefly, cortices from E14.5-E18.5 embryos were dissected in ice-cold 1X HBSS containing 10 mM HEPES and digested in 20 U/ml papain enzyme at 37°C for 10 minutes. 10 mg/ml trypsin inhibitor was used to terminate digestion. Tissue was washed with Basal Media Eagle supplemented with 0.5% L-glutamine, 1% P/S and 5% FBS (BF) twice at room temperature and triturated three to five times. 300.000 cells were plated onto each well of 24-well plate (pre-coated with laminin and poly-L-lysine) in BF. After 2 hours incubation at 37°C, medium was changed into BF containing N-2 and B-27 supplements (Gibco Life Technologies).

### Immunoassays

Neuro2A and HEK293T cells were maintained in EMEM and DMEM, respectively, each were supplemented with 10% FBS, 1% penicillin/streptomycin, 0.5% L-glutamine (complete media). Plasmids were transfected into primary cortical neurons at 2 day *in vitro* (DIV) using Lipofectamine 2000 (Invitrogen), or HEK293T using Lipofectamine 3000 (Invitrogen), according to manufacturer’s protocol. Neuro2A cells were transfected using 1mg/ml Polyethylenimine (PEI) dissolved in sterile water (Polysciences Inc.). For immunoblotting, protein lysates were collected 2 days after transfection either in ice-cold RIPA buffer (0.05 M Tris-HCl, pH 7.5, 0.15 M NaCl, 1% Triton X-100, 1% Na-DOC, 0.1% SDS) supplemented with protease inhibitor or Lysis Buffer (10mM Tris-HCl pH7.6 containing 0.5% SDS, 2%NP40, 150 mM NaCl, 1 mM EDTA) supplemented with 10 mM iodoacetamide and protease inhibitor. For immunoblotting of mice cortical lysates, cortices at different developmental time points (E14, E16, E17, E18, P0, P7, P14, P21, P28 and adult) were collected in 1X cold HBSS containing 10 mM HEPES and tissues were homogenized in ice-cold RIPA buffer supplemented with protease inhibitor with a Dounce homogenizer. Antibodies and dilutions used were as follows: myc (Santa Cruz Biotechnology, cat.no. SC-40, 1:1000); hnRNP C1+C2 (Abcam, cat.no. ab10294, 1:2000); biotin (gift from Dr Timothy J Mitchison, Harvard Medical School, Boston, MA, 1:20,000); Histone H3 (Cell Signaling Technology, cat.no. 9715S, 1:2500); BETA-ACTIN (Abcam, cat.no. ab6276, 1:5000), KIF2A (Abcam, cat.no. ab37005, 1:5000).

For immunofluorescence staining, both Neuro2A cells and primary cortical neurons were transfected with Lipofectamine2000 (Invitrogen) 2 days after plating and fixed with 4% PFA 2 days after transfection. Neuro2A cells were imaged using LEICA DMI8 SP8 CS/DLS confocal microscope and LasX Life Science software. Primary cortical neurons were imaged using Nikon Eclipse 90i confocal microscope and NIS-Elements AR software. For co-localization analysis, quantification was performed using JACoP plugin which calculates the Manders’ Coefficients M1 and M2 and these values provide a direct measure of the quantity of interest independent of signal proportionality ^62-64^. Sholl analysis was performed with the Sholl analysis plug-in in ImageJ software ^19,65^. Antibodies and dilutions used were as follows: myc (Santa Cruz Biotechnology, cat.no. SC-40, 1:1000); GFP (Santa Cruz Biotechnology, cat.no. sc-8334, 1:1000); GFP (Aves Labs, cat.no. GFP-1020, 1:10000); Alpha Tubulin (Cell Signaling Technology, cat.no. 3873S, 1:1000); Beta III Tubulin (Tuj1) (Abcam, cat.no. ab7751, 1:500); KIF2A (Abcam, cat.no. ab37005, 1:5000); Streptavidin Alexa Fluor®488 conjugate (Life Technologies, cat.no. S32354, 1:5000); Nogo C-4 (Santa Cruz Biotechnology, cat.no. sc-271878, 1:500).

### qRT-PCR

Mice cortices at different developmental time points (E14, E16, E18, P0, P7, P14, P21, P28 and adult) were collected in 1X cold HBSS and total RNA was isolated in Trizol (Thermo Scientific). cDNA was synthesized using Transcriptor High-Fidelity cDNA Synthesis Kit (Roche) and qRT-PCR was performed with Luminaris HiGreen qPCR Master Mix (Thermo Scientific) using a CFX Connect Real-Time PCR detection System (Bio-Rad). *Kif2a* mRNA levels were normalized to *Gapdh* mRNA using the 2^−ΔΔCT^ method. The primer sequences were as follows: Gapdh_F(cgacttcaacagcaactcccatcttcc);Gapdh_R (tgggtggtccagggtttcttactcctt);Kif2a.1_F(ccagggtgaaagaattgactg);Kif2a.1_R(gcttcatggaaagtgaac agc);Kif2a.2_F(gcaaatagggtgaaagaattgactg);Kif2a.2_R(tctgggagacagcttcatgg);Kif2a.1_ex5lon g_F(ctccttcacgtagaaaatccaatt);Kif2a.1_ex5long_R(gtctctgatcatacacataatttcgt);Kif2a.3_ex5shor t_F(cagaatgcacgtagaaaatccaa);Kif2a.3_ex5short_R(tctgaagtctctgatcatacacata)

### *In utero* electroporation and image analysis

*In utero* electroporation experiment was performed as previously described ^66,67^ using E14.5 pregnant CD1 mice. Briefly, timed pregnant mice (E14.5) were anesthetized using isoflurane. The uterine horns were exposed by laparotomy and lateral ventricle of each embryo was injected using pulled glass capillaries (Drummond® PCR micropipettes, pulled as 60 µm diameter) with 333 ng/µl NS (control) or shKif2a plasmids combined with 333 ng/µl resKIF2A.1, resKIF2A.2 or resKIF2A.3 and 333 ng/µl pCAGGS_IRES_tdTomato to visualize transfected neurons and 0.1% Fast Green to monitor the injection (All plasmids were prepared using QIAGEN Endofree Plasmid Maxi Kit). Plasmid DNA was electroporated into the neuronal progenitors located in the ventricular zone by 0.5 pulses at 30 V for 400 ms intervals with an electroporater (BTX Harvard Apparatus, ECM830) using 5mm tweezertrodes (BTX Harvard Apparatus). Uterine horns were placed back in the abdominal cavity, and development of embryos was allowed to continue for three days (E17.5). Embryos brains were harvested and fixed with 4% PFA in PBS overnight at 4°C. Brains were sectioned coronally to 50 µm with a cryostat (CM1950, Leica Biosystems). Optical z-series through 50 µm images were captured using LEICA DMI8 SP8 CS/DLS microscope and maximum intensity projections were generated in LasX Life Science software. Images were opened in Adobe Photoshop CC (Adobe Systems) for positioning the ten bins grid for the cell counting. Number of transfected cells were counted for each bin using the cell counter option of ImageJ (NIH) ^19^ and then data were transferred to Excel (Microsoft Office) for further analysis. Bins 1-5 represent cortical plate, bins 6-9 represent subventricular-intermediate zones, and bin 10 represents ventricular zone. Finally, graphs were plotted using GraphPad Prism 5 software.

### Proximity-dependent biotinylation and mass spectrometry analysis (BioID)

Neuro2A cells were transfected with 60 µg BioID constructs using 1mg/ml PEI. The following day, medium was changed with complete EMEM containing 0.05 mM D-biotin (Life Technologies). 2 days after transfection, protein lysates were collected in Lysis Buffer supplemented with 10 mM iodoacetamide and protease inhibitor by lysing the cells using an insulin syringe seven times. Protein concentration was measured by BCA assay (Thermo Scientific) and equal amount of protein from each condition was mixed with equal volume of Streptavidin Plus UltraLink® Resin (Pierce®) which were pre-washed with 1x PBS and Lysis Buffer. Proteins were incubated with streptavidin beads overnight at 4°C on a tube rotator. In order to eliminate sample loss, on-bead digest for streptavidin captured biotinylated proteins was performed the following day. Beads were washed with Wash buffers 1, 2, 3 and 4 two times for 10 minutes each, respectively (Wash buffer 1: 2% SDS; Wash Buffer 2: 2% Na-DOC, 1% Triton X-100, 50 mM NaCl, 50 mM HEPES pH 7.5, 1 mM EDTA; Wash Buffer 3: 0.5% NP-40, 0.5% Na-DOC, 1% Triton X-100, 500 mM NaCl, 1 mM EDTA, 10 mM Tris-HCl pH 8.1; Wash Buffer 4: 50 mM Tris-HCl pH 7.4, 50 mM NaCl). For affinity capture experiment followed by immunoblotting, proteins were eluted with elution buffer (500 nM D-biotin in 2X Laemmli Buffer with 0.1 M DTT) at 98°C, for 10 minutes with 1000 rpm shaking.

For identification of biotinylated proteins by mass spectrometry beads were washed with urea solution (8M urea in 0.1M Tris-HCl pH 8.5) and then reduced with DTT (100 mM in urea solution) at 56°C for 30 minutes for reduction. Beads were washed with urea solution and then incubated with iodoacetamide (0.05 M in urea solution) for 20 minutes under dark for alkylation of cysteine residues. Beads were washed with urea solution and 0.05 M ammonium bicarbonate solution (NH_4_HCO_3_) (0.05 M in HPLC grade water) twice for each buffer. For 600 µg of protein digestion, 4 µg of trypsin (MS Grade Trypsin Protease, Pierce) was added to beads for 16 hours at 37°C with 1000 rpm shaking. Peptides were treated with 10% acetic acid in order to terminate trypsin digestion and peptides were eluted using C18 stage tips ^68^. Eluates were dried in speed vacuum system and were analyzed using a Thermo Scientific Q-Exactive LC-MS/MS mass spectrometer. The data sets were analyzed against the *Mus musculus* Swissprot/Uniprot database ^69^. For analysis, the spectral counts of proteins were used to calculate fold change ratios and FDR (false discovery rate) values for each KIF2A isoform and KIF2A^H321D^ mutant using the qspec-param programme of qprot_v1.3.5. Proteins found in less than two biological replicates were filtered away. Proteins with the same gene name were merged via sum of PSM values. Proteins were filtered with a 0.05 cut-off for Z-statistics and FDR values. Proteins with less than 1.5-fold change are filtered away. Heat map was created with Prism 8. For cluster analysis, significant protein hits were analyzed using the String database v11.0 ^32^ by Cytoscape StringApp ^30^ with 0.4 confidence. Affinity propagation clustering of the network was performed by Cytoscape (version 3.7.2) and its plugin clustermaker ^29^. GO and KEGG enrichment analysis of the network was performed via g:Profiler ^31^.

### Proximity Ligation Assay (PLA)

Neuro2A cells were plated on 24 well plate with coverslips. The proximity ligation assay (PLA) was performed using Duolink® In Situ Red Starter Kit Mouse/Rabbit (DUO92101, Sigma-Aldrich) according to manufacturer’s protocol. Briefly, cells were fixed with 3.2% PFA for 15 min and blocked with Duolink® Blocking Solution for 30 min at 37°C. Cells were incubated with primary antibody pairs or only one primary antibody in Duolink® antibody diluent overnight at 4°C. Cells were washed with Wash Buffer A and incubated with Duolink® Plus and Minus PLA probes in antibody diluent for 1 hour at 37°C. Cells were washed with Wash Buffer A and ligation was performed using ligase diluted in Duolink® Ligation buffer for 30 min at 37°C followed by washes with Wash buffer A. Amplification was performed with polymerase diluted in Duolink® Amplification buffer for 100 min at 37°C. Cells were washed with Wash buffer B followed by 0.01X Wash buffer B and finally mounted with mounting medium containing DAPI. Images were captured with LEICA DMI8 SP8 CS/DLS microscope using DAPI channel to observe nuclei and red channel (λex 594 nm; λem 624 nm) to observe PLA dots. Optical z-series images were projected into a flattened plane with maximum intensity option of the LasX Life Science software. Image analysis was performed with ImageJ (NIH) software ^37^. Antibodies and dilutions used were as follows: KIF2A (Abcam, cat.no. ab37005, 1:5000); RTN4 (Nogo C-4, Santa Cruz Biotechnology, cat.no. sc-271878, 1:500).

### Use of animals

All animal experiments were performed in accordance with the guidelines for the care and use of laboratory animals of Koç University and the Ministries of Food, Agriculture and Livestock, Forestry and Water Management of Turkey. All mice were bred, maintained and housed at the Koç University Animal Facility. Ethics approval was obtained from Koç University Institutional Animal Care and Use Committee (license no. 2014–9).

## Acknowledgements

We thank Ahmet Kocabay, Nilhan Coskun, Mustafa Demir and Muhammed Resat Oguzkan Yaralı (Koç University Animal Research Facility, Turkey) for their help in providing timed pregnant mice, the Molecular Imaging Core Facility of Koç University Research Center for Translational Medicine (KUTTAM), Berna Morova and Alper Kiraz from Koç University College of Science for assistance with confocal microscopy, and Cory Dunn for feedback on the manuscript. The work was funded by following grants to G. Ince-Dunn: European Commission FP7 International Reintegration Grant (PIRG07-GA-2010-268433), Scientific and Technological Research Council of Turkey (TUBITAK-151Z707) and to C. Akkaya: TUBITAK-BIDEB 2211/E Scholarship Program.

## Author Contributions

C.A. carried out all cell and molecular biology experiments, *in utero* migration and BioID assays, analyzed the data and wrote the manuscript. D.A., E.B. and A.C.T. assisted with *in utero* electroporations. A.K. carried out gene ontology analysis and interactome mapping. B.A.A. performed mass spectrometry. G.G. cloned Kif2a isoforms. N.O. supervised the BioID assay and mass spectrometry experiments and analyzed data. G.I.D. conceived the hypothesis, designed research, analyzed data and wrote the paper. All authors discussed the results and reviewed the manuscript.

---

**Supplementary Figure 1.**
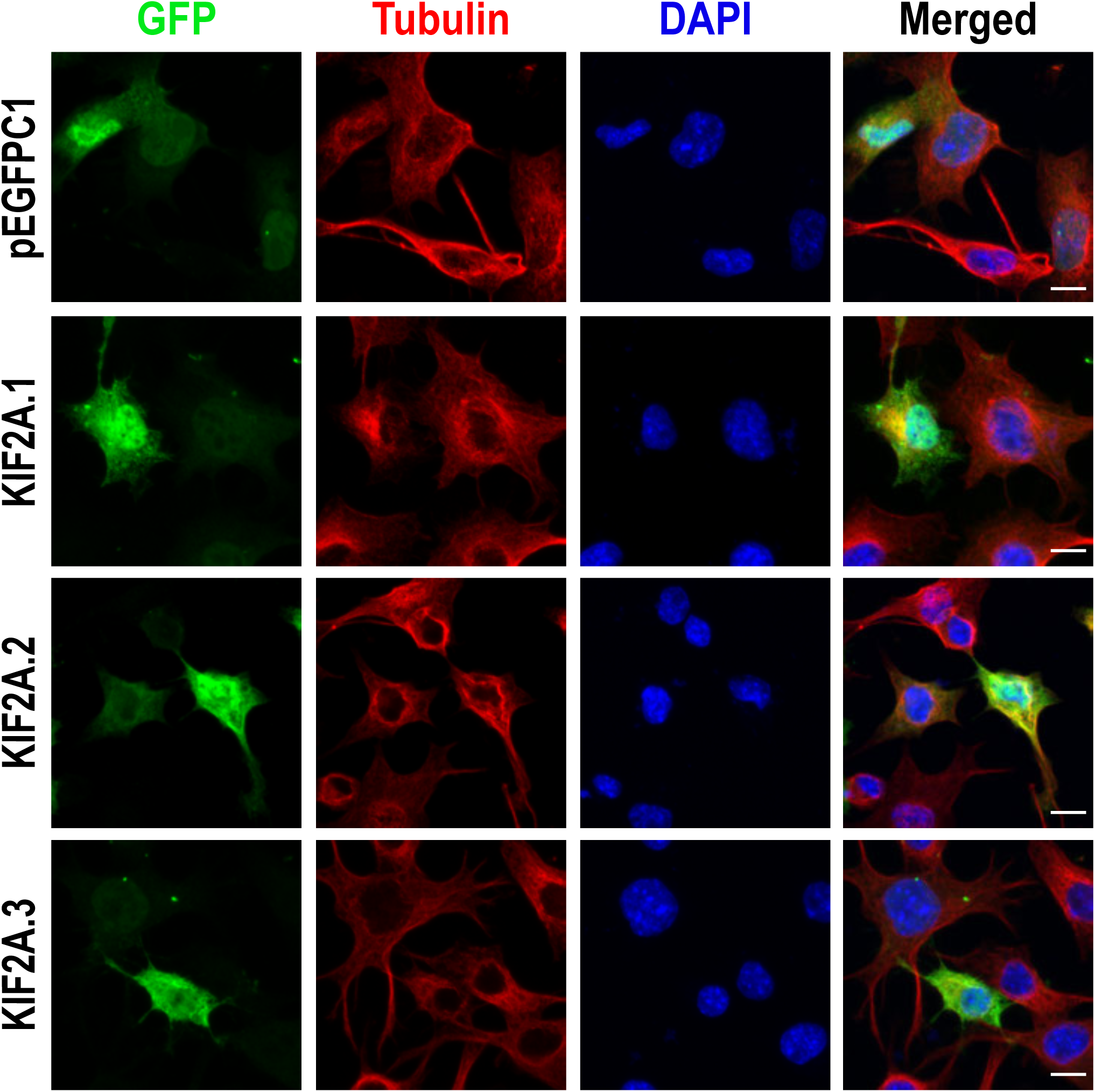
All KIF2A isoforms have diffused localization to cytoplasm and nucleus in Neuro2A cells. Neuro2A cells transfected with plasmids expressing pEGFPC1-tagged KIF2A isoforms (KIF2A.1, KIF2A.2 and KIF2A.3) 2 days after plating, fixed with 4% PFA 2 days after transfection and immunostained with anti-GFP (green). Anti-αTubulin visualized microtubules (red) and DAPI stained DNA. Scale bar, 10 µm.

**Supplementary Figure 2.**
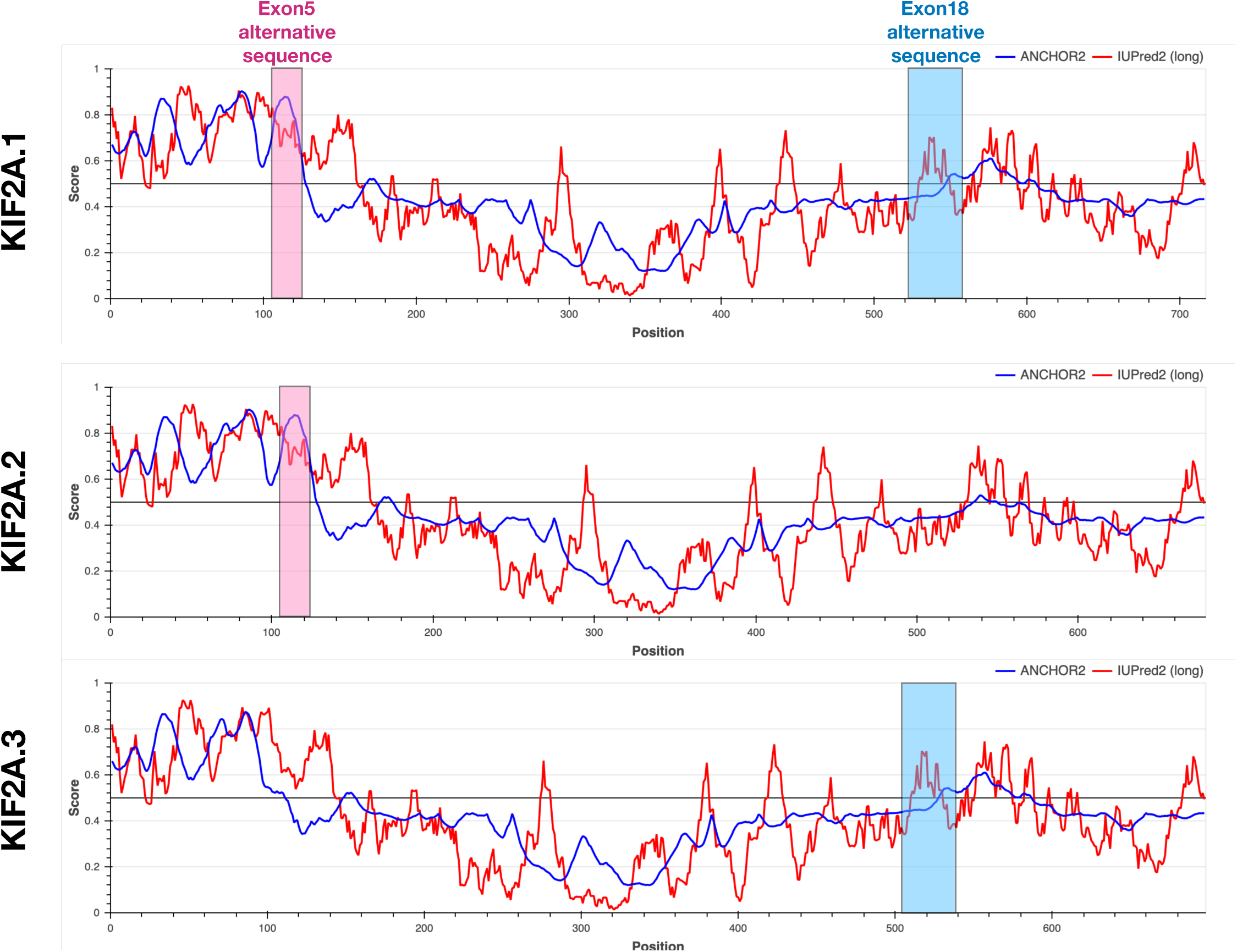
Measure of disorder in protein regions of individual KIF2A isoforms. A measure of protein disorder is plotted as a function of amino acid sequence of individual KIF2A isoforms. The IUPred2 and ANCHOR2 algorithms are used ^1^. Alternative sequences are labeled in pink (exon5) and blue (exon18).

**Supplementary Figure 3.**
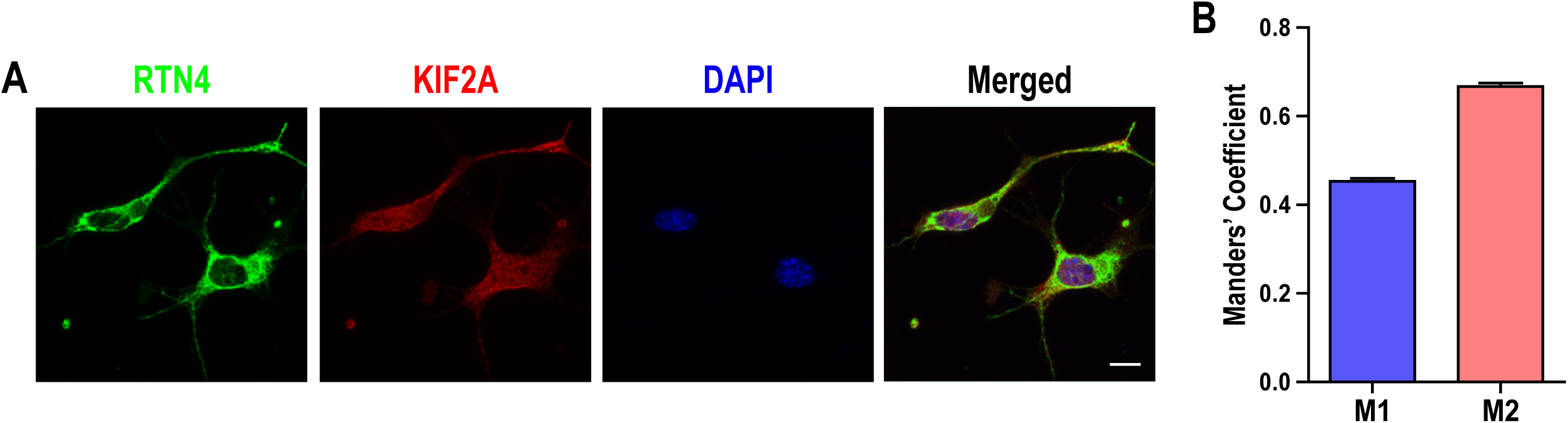
KIF2A and RTN4 are partially co-localized in Neuro2A cells. **A**. Immunofluorescence staining of RTN4 (green), KIF2A (red), DNA (blue) in Neuro2A cells. Scale bar, 10µm. **B**. Quantification of co-localization of KIF2A with RTN4A using Manders’ coefficient ^2-4^. M1: Fraction of KIF2A fluorescence overlapped with RTN4 fluorescence. M2: Fraction of RNT4 fluorescence overlapped with KIF2A fluorescence. n=219, data is represented as bar graphs. Bars represent S.E.M.

**Supplementary Figure 4.**
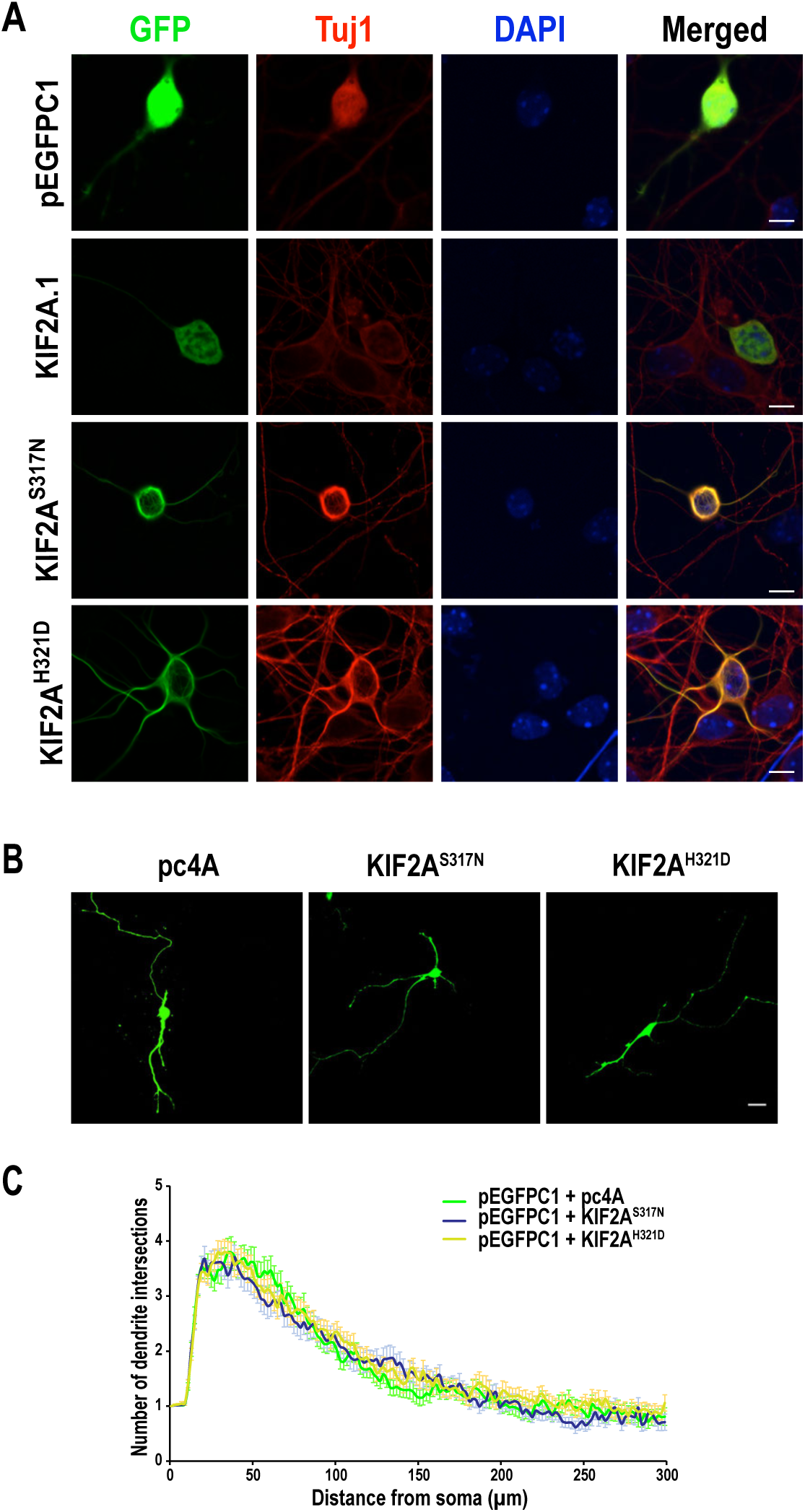
*Kif2a* disease mutants co-localize predominantly to microtubules and do not affect dendrite development in primary cortical neurons. **A**. E18.5 primary cortical neurons were transfected with plasmids expressing pEGFPC1-tagged KIF2A.1, KIF2A^S317N^ or KIF2A^H321D^ and immunostained with anti-GFP (green) and anti-Tuj1 (red). DNA was stained with DAPI. **B**. Representative images of E14.5 primary cortical neurons co-transfected with pEGFPC1 along with *Kif2a* mutants. **C**. Dendrite development was quantified by Sholl analysis ^5,6^. n=70 for each condition derived from three separate neuronal cultures. Unpaired *t* test determined the p value. p<0.05. Scale bar in A and B, 10µm.

## Notes

### Competing Interest Statement

The authors have declared no competing interest.

